# Ligand binding characteristics of an NAD+ riboswitch revealed by FRET and biolayer interferometry

**DOI:** 10.1101/2024.11.10.622884

**Authors:** Nico E. Conoan Nieves, Julia R. Widom

**Affiliations:** University of Oregon Department of Chemistry and Biochemistry

## Abstract

The Class II NAD^+^ riboswitch is a bacterial RNA that binds ligands containing nicotinamide. Herein, we report a fluorescence and biolayer interferometry study of riboswitch interactions with β-NMN. The results reveal a shift in the prevalence of a pseudoknot structure in the presence of ligand and Mg^2+^.

---

Riboswitches are a topic of increasing interest due to their potential as antibiotic targets^1^ and molecular sensors^2^. A riboswitch is a gene regulatory element, commonly found in the 5” untranslated region of bacterial messenger RNA, that contains an aptamer domain where a ligand binds and an expression platform located just downstream. The Class II NAD^+^ riboswitch (NAD^+^-II) is a translational off-switch that has been reported to bind specifically to the nicotinamide moiety of nicotinamide adenine dinucleotide (NAD^+^) (Fig. 1a), a ubiquitous cellular cofactor involved in many enzymatic reactions.^3^ We aimed to elucidate the conformational dynamics and ligand-binding interactions of NAD^+^-II using fluorescence techniques including single-molecule Förster resonance energy transfer (smFRET). This method relies on a donor and acceptor pair of fluorophores, selected such that an excited donor fluorophore may nonradiatively transfer energy to the acceptor fluorophore with an efficiency that depends on their proximity and relative transition dipole orientations.^4,5^ FRET is sensitive to changes in inter-fluorophore distance on the nanometer length-scale, and modern single-molecule instruments can routinely resolve changes in FRET on the tens-of-milliseconds timescale.^6^ Appropriate fluorophore labeling sites will change distance as the RNA samples different conformations without interfering with the folding mechanism or binding interaction under investigation.^7^

**Fig. 1.**
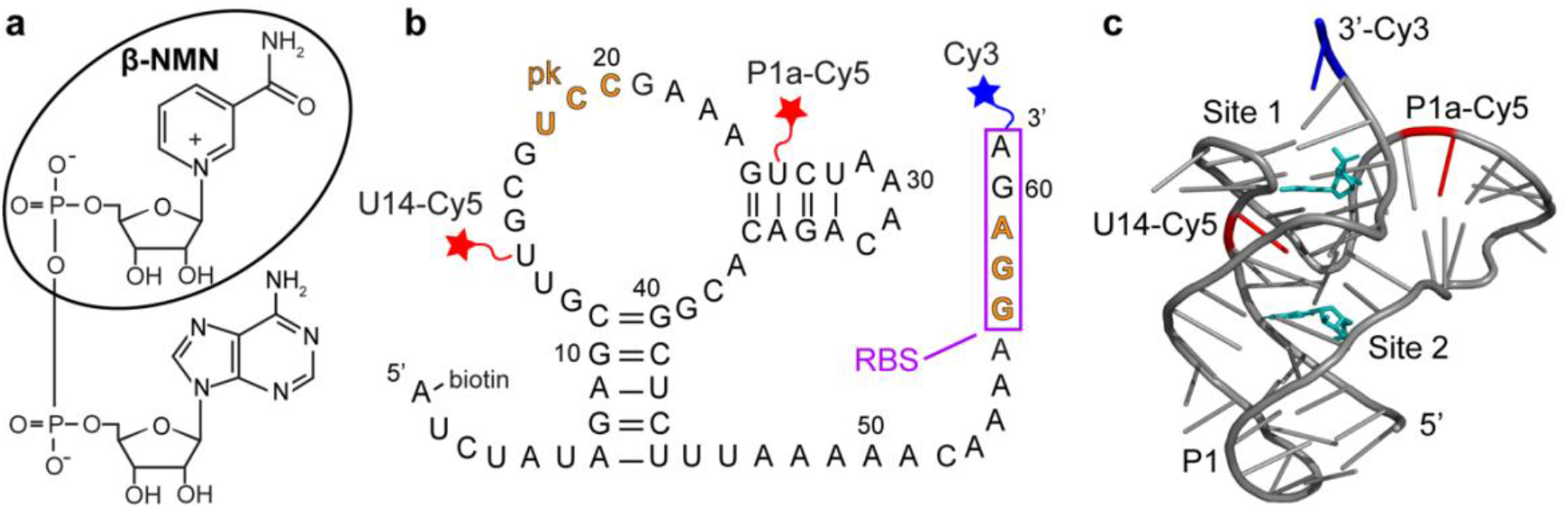
(a) Chemical structure of nicotinamide adenine dinucleotide (NAD^+^). The circled moiety is β-nicotinamide mononucleotide (β-NMN). (b) Sequence and secondary structure of the NAD^+^-II riboswitch from *S. parasanguinis*. The ribosome binding site (RBS) is indicated by a box and the bases that form the pseudoknot (pk) are indicated in orange. Cy3 (blue star) is placed at the 3” end, and Cy5 (red stars) is placed at either U14 (b-U14Cy5-Cy3 riboswitch construct) or U26 (b-P1aCy5-Cy3 construct). (c) Crystal structure of the riboswitch (PDB ID 8HB8) with the P1 and P1a domains and β-NMN binding sites labeled. Locations of the U14 and P1a Cy5 labeling sites are indicated in red and β-NMN molecules are shown in cyan.

A biochemical study of NAD^+^-II reported the potential for the formation of a pseudoknot in which three bases within the ribosome binding site (RBS) form base pairs with three bases within a loop near the 5” end of the aptamer domain (Fig. 1b).^3^ Titrations based on in-line probing^3^ and calorimetry^8^ reported a 1:1 binding stoichiometry with KD values of ∼100 µM for NAD^+^ and ∼40 µM for β-nicotinamide mononucleotide (β-NMN). Two crystallography studies further reported NAD^+^- and NMN-bound structures, finding that the primary binding pocket of the riboswitch is specifically sensitive to the nicotinamide moiety.^8,9^ One study observed RNA dimerization with each monomer binding one ligand molecule^8^ and the other observed RNA monomers, each of which binds two ligands in distinct domains^9^ (Fig. 1c). The overall ligand-bound RNA structures in both reports are in good agreement with each other and both demonstrate RBS sequestration through pseudoknot formation, supporting the translational off-switch mechanism originally proposed for this riboswitch.^3^

We designed two riboswitch variants based on the *pnuC* motif from *S. parasanguinis*, labeled with Cy3 and Cy5 as the FRET donor and acceptor, respectively. In both cases, the 5” end of the RNA is biotinylated for surface immobilization, Cy3 is located on the 3” end, and Cy5 is placed internally via conjugation to a 5-aminohexylacrylamino-uridine modification. Labeling sites were selected based on those previously used for the pseudoknot-structured Class I preQ1 riboswitch.^10,11,12^ Specifically, Cy5 labeling positions of U14 (in the loop adjacent to the P1 helix) and U26 (in the P1a helix) were selected to ideally result in a high-FRET (HF) state when the riboswitch exists in its compact “docked” pseudoknot structure and a mid-FRET (MF) or low-FRET (LF) state when undocked. We refer to these constructs as b-U14Cy5-Cy3 (abbreviated U14-Cy5) and b-P1aCy5-Cy3 (abbreviated P1a-Cy5) RNAs based on the locations of the 5” biotin (b), internal Cy5, and 3” Cy3 (Fig. 1b). The acrylamino-modified U14 construct prior to Cy5 labeling is referred to as b-aminoU14-Cy3 (abbreviated aminoU14).

We first performed smFRET on the U14-Cy5 riboswitch in the absence and presence of 1 mM Mg^2+^, as divalent cations support RNA folding by coordinating to its 2” hydroxyl groups and negatively charged phosphate backbone. Cy3 and Cy5 fluorescence were recorded in real-time and segments of time traces indicating activity of both fluorophores were manually selected using custom MATLAB code. After acquiring >200 traces, the first 50 frames of each were compiled to populate histograms that display the relative frequencies of the FRET efficiencies accessed by the RNA. Three FRET states are observed in the histogram for apo-form RNA in the absence of Mg^2+^ with FRET efficiencies of EHF = 0.92, EMF = 0.64 and ELF = 0.16, with the HF state being the most prevalent (Fig. 2a). The prevalence of the MF state increases slightly in the presence of 1 mM Mg^2+^ (Fig. S2). After increasing the divalent cation concentration to 20 mM Mg^2+^, the concentration used most widely in solution-phase studies,^3,8^ the MF state is substantially suppressed in favor of the HF state (Fig. 2a). With the fluorophores placed adjacent to the two arms of the helix forming the pseudoknot, we hypothesized that addition of ligand would further shift population to the HF state. Surprisingly, upon introducing 1 mM β-NMN in combination with either 1 mM or 20 mM Mg^2+^, no significant difference is observed in histograms or time traces compared to the same Mg^2+^ concentration in the absence of ligand. This observation could be explained by the riboswitch preferring the pseudoknot structure, or a different HF structure, under all conditions tested. Alternatively, fluorophore placement may either not be sensitive to the particular structural changes undertaken by the RNA or may be impairing the ability of the riboswitch to bind ligand altogether.

**Fig. 2.**
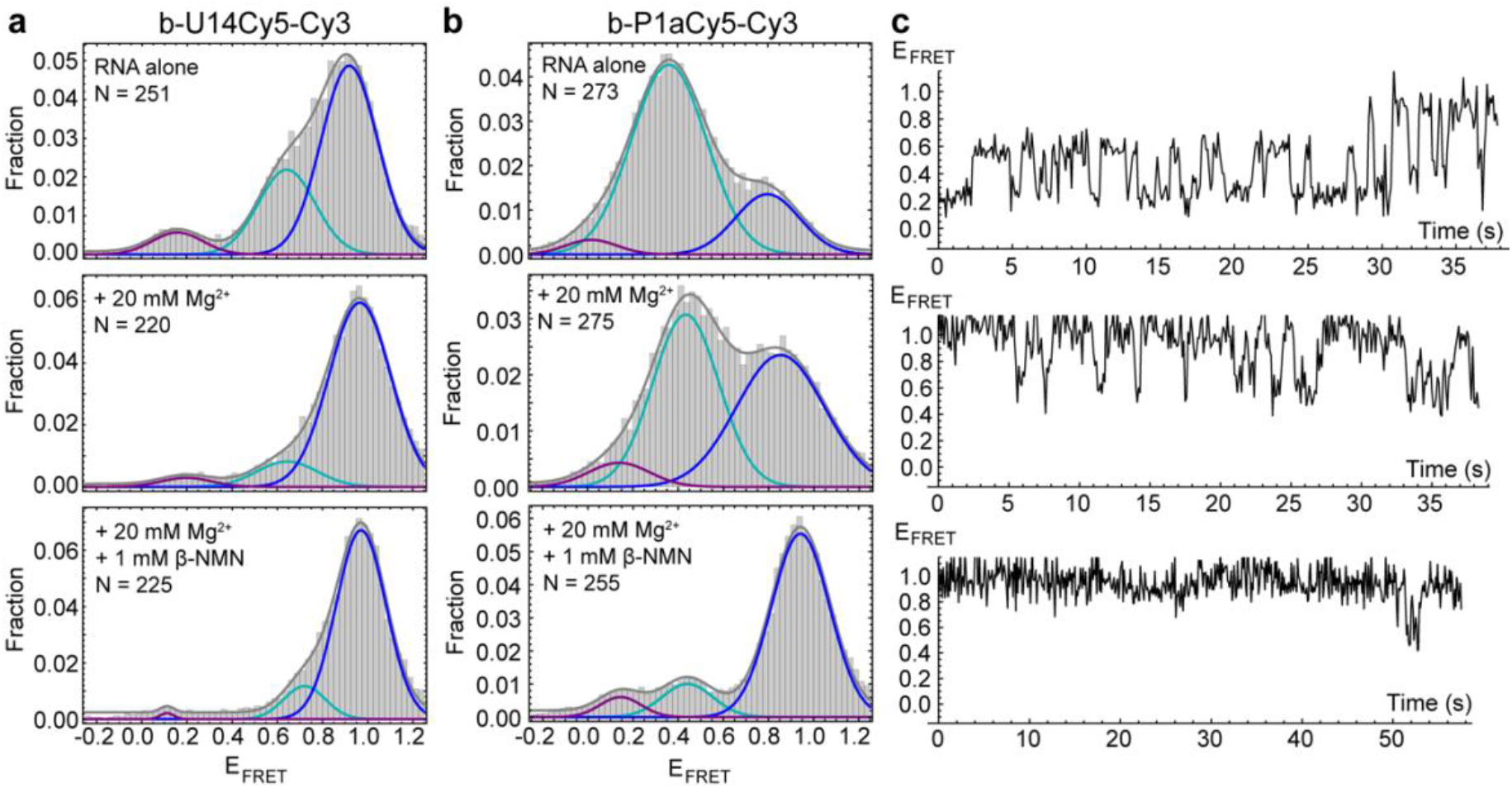
(a) smFRET histograms of the U14-Cy5 riboswitch construct recorded in imaging buffer (top), buffer+20 mM MgCl_2_ (middle) or buffer+20 mM MgCl_2_+1 mM β-NMN (bottom). 3-Gaussian fits are shown in gray, with underlying components shown in purple, cyan and blue. “N” indicates the number of single-molecule traces that were used to compile the histogram. (b) Corresponding smFRET histograms of the P1a-Cy5 construct. (c) Representative FRET efficiency traces from single molecules of P1a-Cy5 RNA in imaging buffer (top), buffer+20 mM MgCl_2_ (middle) and buffer+20 mM MgCl_2_+1 mM β-NMN (bottom)

Inspection of the crystal structures indicated that placement of Cy5 on U14 has the potential to interfere with ligand binding. The nicotinamide moiety of NAD^+^ hydrogen-bonds to form a planar “base” triple and intercalates into a four-rung ladder of base triples and quadruples.^8,9^ U14 is immediately adjacent to the base quadruple that forms a “lid” on the nicotinamide binding pocket (Fig. 1c). We utilized biolayer interferometry (BLI) to directly investigate whether ligand was binding to the U14-Cy5 riboswitch using the aminoU14 RNA, which lacks Cy5, as a control. The aminoU14 RNA was observed to bind β-NMN, whereas the U14-Cy5 RNA did not respond when exposed to 1 mM β-NMN in the presence of 20 mM Mg^2+^ (Fig. 3a). These results indicate that the Cy5 modification suppresses ligand binding, either by directly blocking or altering the binding sites, or by promoting misfolding of the RNA. Fitting the BLI trace of aminoU14 RNA revealed fast and slow components to the association phase, which is most readily explained by a 2:1 binding stoichiometry (as seen in one crystal structure^9^) with ligand accumulating more slowly at one binding site than the other (Fig. 3b). The slow rate at which a second binding site appears to get populated may explain why solution-phase measurements have typically suggested a 1:1 binding stoichiometry.^3,8^

**Fig. 3.**
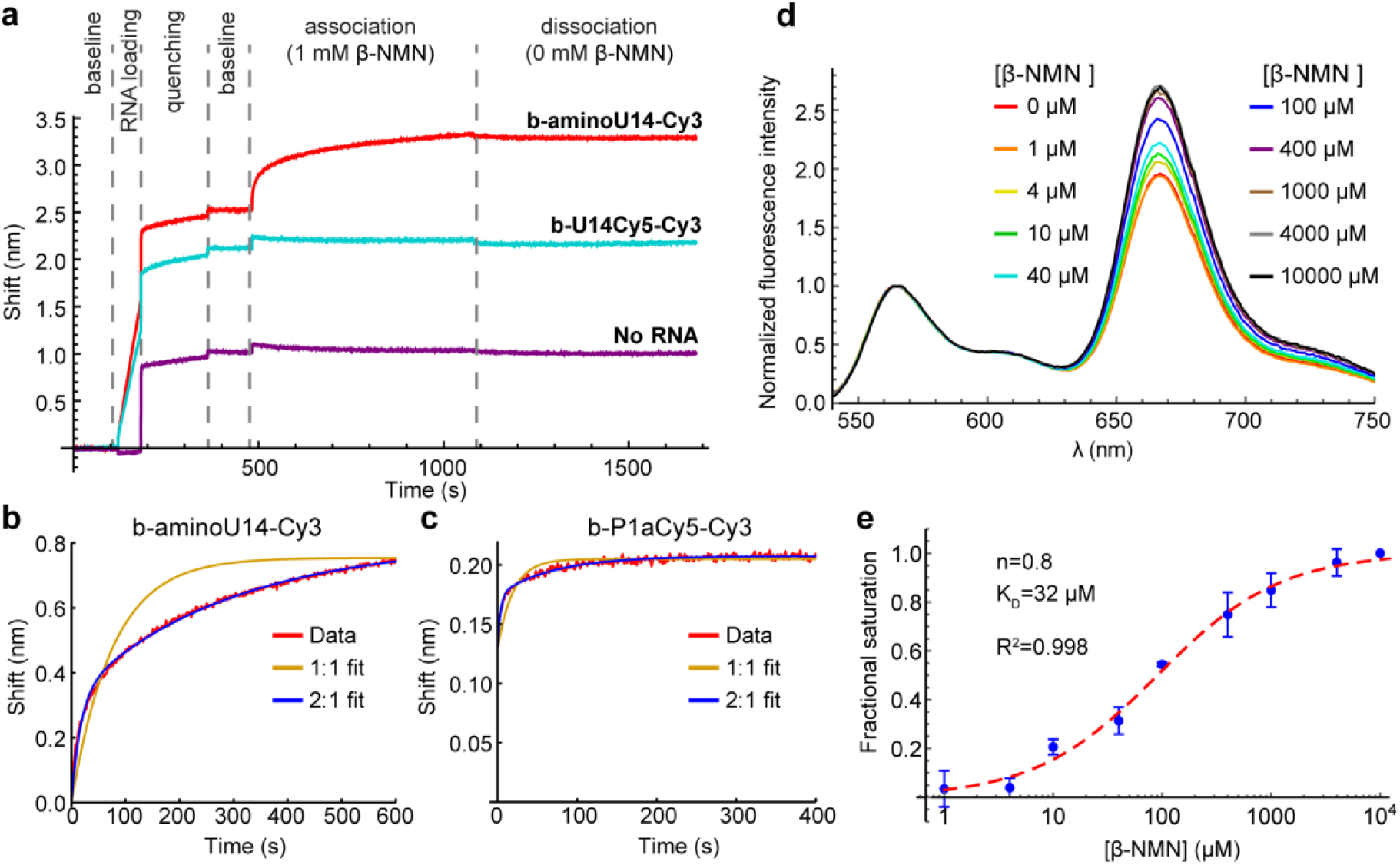
(a) Biolayer interferometry (BLI) measurements on aminoU14 (red) and U14-Cy5 RNA (cyan). Shifts are observed due to binding of RNA to the streptavidin-coated probe (“RNA loading”), binding of biocytin to the probe (“quenching”) and binding of β-NMN to the RNA (“association”). A probe was run with no RNA loaded to control for nonspecific binding of β-NMN to the probe (purple). (b) Close-up of the association phase for aminoU14 RNA (red), showing fits for a 1:1 (yellow) and 2:1 (blue) β-NMN:RNA binding stoichiometry. (c) Close-up of the association phase for P1a-Cy5 RNA and fits. (d) Bulk FRET titration of P1a-Cy5 RNA with β-NMN. Emission spectra were recorded under excitation at 532 nm and the Cy3 peak was normalized to an intensity of 1. (e) Blue: titration curve extracted from the ratio of Cy5 to Cy3 emission intensity. Datapoints show the mean and standard deviation across three titrations. The curve was fit with the Hill equation (red dashed line), yielding a cooperativity of 0.8 and a K_D_ of 32 µM.

We identified an alternative Cy5 labeling site in the P1a helix (U26), which exhibits the highest sequence variability in the aptamer domain and does not appear to be directly involved in ligand binding.^3,8,9^ This segment of P1a resides near the 3” end of the RNA in crystal structures, suggesting that a HF state would be favored in the presence of ligand. BLI results confirm functionality and again show fast and slow components to the association phase when the P1a-Cy5 riboswitch is exposed to β-NMN (Fig. 3c and S3). Bulk FRET titrations over the range of 0-10 mM β-NMN indicate that there is a global shift to higher FRET efficiencies with increasing ligand concentration (Fig. 3d), which was not seen in a mock titration (only buffer added) or a negative control titration with the related compound nicotinic acid mononucleotide (NaMN; Fig. S4). Fitting the ratio of Cy5 to Cy3 intensity with the Hill equation after correction for photobleaching (details in methods) yielded a *KD* of 32 µM and a Hill coefficient, a reporter on the cooperativity of binding sites, of *n* = 0.8 (Fig. 3e). This *K*_*D*_ is comparable to previously reported affinities for β-NMN^3,8^, confirming that the modifications on the P1a-Cy5 riboswitch do not significantly hinder ligand binding.

smFRET measurements revealed that the P1a-Cy5 riboswitch briefly samples several distinct conformations in the absence of Mg^2+^ and ligand, which are observable as two prominent features in FRET histograms. A broad MF peak centered on E_FRET_ = 0.36 is dominant (Fig. 2b) and inspection of traces suggests that several different FRET states may be resolvable within this peak (Fig. 2c). The riboswitch”s dynamics are substantially suppressed upon adding 20 mM Mg^2+^, with HF states (E_FRET_ > 0.8) being stabilized relative to MF states. In the presence of 1 mM β-NMN and 20 mM Mg^2+^, the complexity of the traces further simplifies to three commonly accessed states with ELF = 0.14, EMF = 0.44, and EHF = 0.94. The prevalence of the HF state increases significantly, and a greater fraction of traces exhibit static rather than dynamic behavior. This response to ligand is reminiscent of the Class I preQ1 riboswitch, which like NAD^+^-II has a pseudoknot-structured architecture and adenosine-rich 3” tail.^10,11,12^ Overall, our smFRET data indicate that the ensemble of structures sampled by the NAD^+^-II riboswitch is more complex than simply undocked or docked (pseudoknot-structured) conformations and that the dynamic sampling of these structures is suppressed in the presence of ligand.

We next tested the potential of the riboswitch to form dimers (as observed in one crystal structure^8^) in solution. Nondenaturing polyacrylamide gel electrophoresis did not indicate the presence of stable dimers, even at high RNA concentrations (Fig. 4a and S5). To further probe the potential for dimer formation, we performed a single-molecule colocalization study using two single fluorophore-labeled riboswitches: the biotinylated aminoU14 RNA discussed above (b-aminoU14-Cy3) and a 3” Cy5-labeled RNA. Several scenarios were tested to provide different opportunities for homo- and heterodimers to form and be observed, including different annealing and imaging conditions (Fig. 4b-e and S6). Homodimers were observed as multi-step photobleaching of Cy3 (Fig. 4c). Transient or extended observations of Cy5 revealed the presence of heterodimers (Fig. 4d-e).

**Fig. 4.**
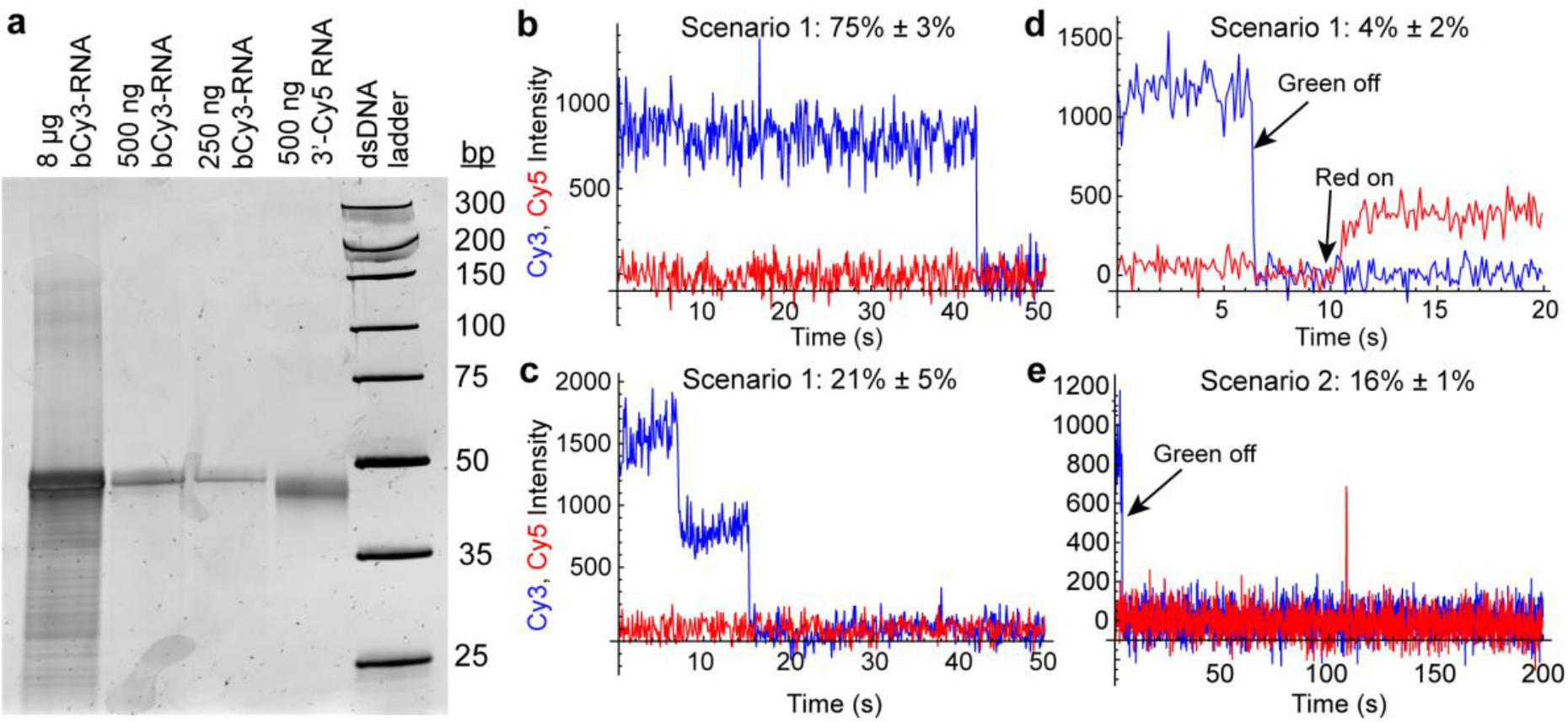
(a) Native polyacrylamide gel showing the presence only of a monomer band (61 nucleotides) when varying quantities of b-aminoU14-Cy3 RNA (lanes 1-3) or 3”-Cy5 RNA (lane 4) are loaded. Weak slower-migrating bands are observed only in an overloaded lane. (b) Example of a trace showing single-step photobleaching of Cy3 on a slide prepared under dimerization scenario 1 (see Fig. S4). (c) Example of a trace showing two-step photobleaching of Cy3 and no Cy5 obtained from a slide prepared under dimerization scenario 1. (d) Example of a trace showing a stable heterodimer on a slide prepared under dimerization scenario 1. The green laser was turned off at 6 s and the red laser was turned on at 10 s. (c) Example of a trace showing Cy3 and transient presence of Cy5 obtained from a slide prepared under dimerization scenario 2 (see Fig. S4). The green laser was turned off at 4 s and the red laser was on continuously.

When amino-U14 and 3” Cy5 RNAs were annealed in a 1:10 ratio before immobilization (scenario 1, Fig. S6a), we observed a small fraction of traces exhibiting Cy5 signal under direct excitation, 4% ± 2% (mean ± standard deviation, σ, across 4 movies). When b-aminoU14-Cy3 RNA was annealed alone, followed by imaging with freely diffusing 3” Cy5-RNA present in the sample chamber (scenario 2, Fig. S4b), transient heterodimer observations occurred in 16% ± 1% (mean ± σ across 2 movies) of traces (Fig. 4e). Multi-step photobleaching of Cy3 (Fig. 4c), observed in 21% ± 5% (mean ± σ across 3 movies) of traces in scenario 1, can result from dimerization or from independent immobilization of two molecules within the diffraction-limited width of the point-spread function.^13^ The annealing conditions in Scenario 1 were designed to favor the formation of heterodimers over homodimers, so it is likely that many of these Cy3 colocalizations are coincidental. Therefore, our data do not support the extensive formation of stable RNA dimers in solution in the absence of Mg^2+^ and ligand.

In summary, we report an investigation into the binding interactions of the NAD^+^-II riboswitch and β-NMN. Our results indicate that the riboswitch dynamically samples a complex landscape of conformations with compact structures being favored in the presence of Mg^2+^ and static occupation of a compact conformation being favored in the presence of β-NMN. We found that stable RNA dimers are not prevalent in solution in the absence of Mg^2+^ and ligand, and our BLI measurements suggest a 2:1 binding stoichiometry of ligand to RNA. Our findings also emphasize the potential impacts of fluorophore labeling on riboswitch function, with ligand binding being inhibited when Cy5 is conjugated to U14. Ongoing work includes a more extensive smFRET study of the NAD^+^-II riboswitch in which we further investigate its dynamics as well as its interactions with NAD^+^ and other ligand variants.

## Author contributions

N.E.C.N.: Conceptualization, Investigation, Visualization, Writing - original draft. J.R.W.: Conceptualization, Supervision, Funding Acquisition, Visualization, Writing - revision and editing.

The authors thank the National Institutes of Health (R35-GM147229) for supporting this research. We also thank the labs of Marian Hettiaratchi and Parisa Hosseinzadeh at the University of Oregon for the use of their biolayer interferometry instrument and Justin Svendsen for training and guidance.

## Data availability

Data for this article, including single-molecule histogram files, fluorescence spectra, gel electrophoresis images and biolayer interferometry traces, are available through Dryad at DOI: 10.5061/dryad.4b8gthtnw

