## Supplemental information for "Ligand binding characteristics of an NAD+ riboswitch revealed by FRET and biolayer interferometry"

Table of Contents

1. Materials and Methods
2. Supplementary Figures
    a. Figure S1: UV/vis spectrum of riboswitch
    b. Figure S2: smFRET in 1 mM $MgCl_2$
    c. Figure S3: Complete BLI traces of P1a-modified riboswitch
    d. Figure S4: Raw fluorescence spectra from bulk FRET titrations
    e. Figure S5: Schematics of dimerization scenarios
    f. Figure S6: Raw gel image
3. Table of RNA Sequences

### 1. Materials and Methods

Materials: RNA was obtained from Horizon Discovery and HPLC-purified and deprotected by the manufacturer. RNA concentrations were determined via A260 using the extinction coefficient provided by the manufacturer. Cy5-labeled constructs were labeled using Cy5 mono-reactive dye packs (Cytiva PA25001). One tube of dye was dissolved in 20 μL of anhydrous DMSO and combined with 30 μL of 167 mM $Na_2CO_3$ containing 3-4 nanomoles of RNA. The reaction was incubated at RT for 2 hours and excess dye was removed via ethanol precipitation. The RNA was then purified via denaturing polyacrylamide gel electrophoresis and recovered via electrocution and ethanol precipitation. RNA was annealed before each experiment by heating to 90 °C for 2 minutes in imaging buffer (50 mM Tris-HCl pH 8.0, 100 mM KCl) and then air-cooling over 10 minutes. $MgCl_2$, β-NMN (Millipore Sigma N3501) and/or NaMN (Millipore Sigma N7764) were added to the desired concentrations after annealing. For single-molecule measurements, the buffer additionally contained an oxygen scavenging system consisting of 50 nM protocatechuate 3,4-dioxygenase (PCD, Millipore Sigma P8279), 5 mM protocatechuic acid (PCA, Millipore Sigma 08992) and 5 mM Trolox (Millipore Sigma 238813). Microscope slides were prepared by the DDS Tween-20 method[1], in which slides are passivated with dichlorodimethylsilane (Millipore Sigma 440272), followed by incubation with biotinylated bovine serum albumin (Thermo Scientific PI29130), streptavidin (Molecular Probes S888) and Tween-20 (Millipore Sigma P9416).

smFRET measurements: Data were collected on a home-built prism-type total internal reflection fluorescence (TIRF) microscope (Leica DMi8 inverted microscope, 63x oil immersion objective). Cy3 and Cy5 were excited by a 532 nm laser (Coherent OBIS 532 LS) at 100-120 mW and a 637 nm laser (Coherent OBIS 637 LX) at 60-75 mW, respectively. The slide was illuminated at 532 nm continuously and at 637 nm at the beginning and end of each movie to confirm that Cy5 was active. For U14-Cy5 RNA and dimer study movies, fluorescence was recorded on a Hamamatsu C13440 Orca Flash 4.0 sCMOS with an exposure time of 100 ms. P1a-Cy5 RNA movies were recorded on an Andor iXon 888 emCCD with an exposure time of 40 ms. Traces exhibiting single-step photobleaching of Cy3 and Cy5 were selected manually and analyzed using custom MATLAB codes. Histograms were compiled from the first 50 frames of each selected trace. Gaussian fitting of histograms was performed in Mathematica.

Bulk FRET measurements: Fluorescence spectra for bulk FRET measurements were collected using an Edinburgh Instruments FS5 fluorometer by exciting Cy3 at 532 nm and recording emission from 540 nm to 750 nm. β-NMN or NaMN was titrated over a range of 0-10 mM into 150 nM RNA in imaging buffer containing 20 mM $Mg^{2+}$. Decreasing fluorescence intensity for both dyes was observed over time due to photobleaching of molecules during each 210-second scan (Fig. S4). To correct for this, we performed a "mock" titration in which in which aliquots of buffer without ligand were added, keeping all other measurement parameters and timing the same. The ratio of the peak intensity of the Cy5 peak to the Cy3 peak was calculated for each spectrum in the β-NMN and mock titrations, and the ratio observed in the mock titration's spectrum was subtracted from the ratio observed in the corresponding β-NMN titration's spectrum (Fig. S3g). This quantity was then converted to fractional saturation (FS) using the following equation:

$$FS(c) = \frac{R_c - R_0}{R_f - R_0}$$

where $R_c$ is the corrected Cy5/Cy3 peak ratio at β-NMN concentration *c*, $R_0$ is the peak ratio at 0 μM β-NMN, and $R_f$ is the peak ratio at the final β-NMN concentration measured, 10 mM. The fractional saturation curve was then fit with the Hill equation:

$$FS(c) = \frac{c^n}{c^n + K_D}$$

where *n* is the Hill coefficient and $K_D$ is the dissociation constant.

Native polyacrylamide gel electrophoresis: After annealing, samples were mixed 1:1 with 2x TBE (Tris-Borate-EDTA) buffer containing 40% glycerol, then run out on a 12% non-denaturing polyacrylamide gel in TBE buffer at 4ºC. The gel was stained in Sybr Gold and imaged on a Typhoon scanner.

Dimer study: In one experimental design ("Scenario 1"), a mixture of b-aminoU14-Cy3 RNA and 3' Cy5-RNA was annealed at a 1:10 ratio in imaging buffer. The mixture was flowed onto the microscope slide, which was then flushed with buffer prior to imaging. Annealing at this ratio would allow for preferential formation of heterodimers over homodimers. To observe monomers and homodimers, the samples were excited at 532 nm until photobleaching of Cy3 occurred. Traces were categorized by whether they exhibited single- or double-step blinking and photobleaching. Spots that exhibited two-step blinking or photobleaching indicate either homodimers or independent colocalized molecules. Spots exhibiting Cy5 signal under excitation with a red laser were classified as heterodimers. In another design ("Scenario 2"), the RNAs were annealed separately, b-aminoU14-Cy3 RNA was immobilized onto the slide, then 100 pM 3' Cy5 RNA was introduced and allowed to freely diffuse while movies were collected. The slide was illuminated briefly at 532 nm to localize the b-aminoU14-Cy3 molecules, then illuminated at 637 nm to view Cy5 colocalization under direct excitation.

Biolayer interferometry: BLI measurements were performed on a GatorPlus Biolayer Interferometer (GatorBio, Paolo Alto, CA) utilizing proprietary GatorOne software for data collection, reference subtraction, and calculation of binding stoichiometry. GatorBio streptavidin-coated glass probes were presoaked for 3 min in imaging buffer containing 1 or 20 mM $MgCl_2$ before collecting a baseline signal for 180 s. The probes were immersed in 100 nM biotinylated RNA until an adequate shift (~1 nm) in signal was observed (1-1.5 min). Probes were then submerged into a 1 ppt biocytin solution to saturate all remaining streptavidin sites. Loaded probes were submerged into imaging buffer to ensure the baseline wavelength reading had stabilized (180 s). To measure association, loaded probes were submerged into 1 mM β-NMN in imaging buffer for 400-600 s. Traces from reference probes with no RNA loaded were subtracted from sample results to correct for non-specific background signal prior to calculating association stoichiometry.

1. Hua, K. Y. Han, R. Zhou, H. Kim, X. Shi, S. C. Abeysirigunawardena, A. Jain, D. Singh, V. Aggarwal, S. A. Woodson, and T. Ha, *Nat. Methods*, 2014, **11**, 1233-1236.

### 2. Supplementary Figures

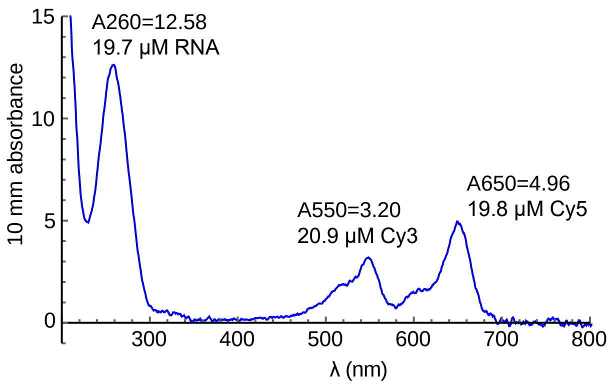


Figure S1. Example UV/visible absorbance spectrum of b-P1aCy5-Cy3 RNA recorded on a Nanodrop spectrophotometer. A260, A550 and A650 (values indicated on the plot) reflect the expected 1:1:1 RNA:Cy3:Cy5 labeling stoichiometry.

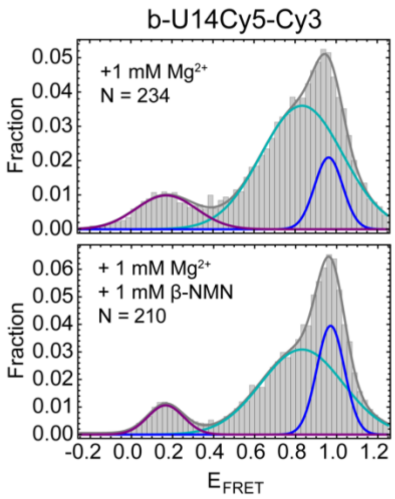


Figure S2. smFRET histograms and 3-Gaussian fits of b-U14Cy5-Cy3 RNA in the presence of 1 mM $MgCl_2$ (top) and 1 mM $MgCl_2$ + 1 mM β-NMN (bottom). “N” indicates the number of single-molecule traces that were used to compile the histogram.

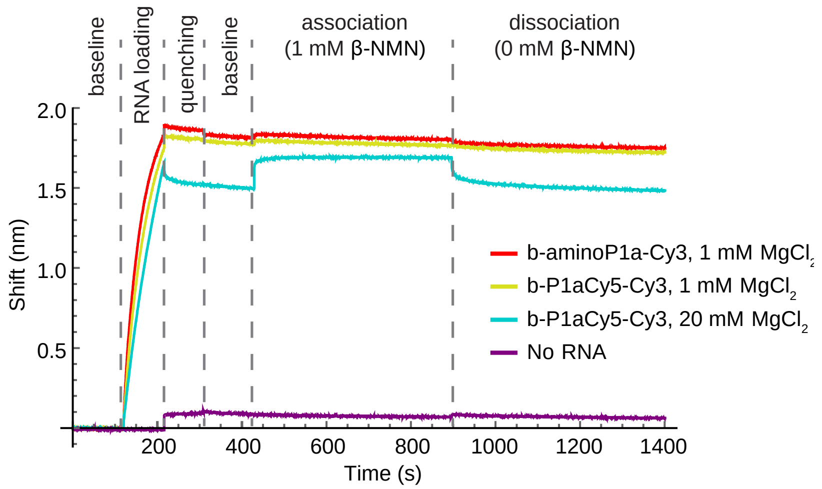


Figure S3. Complete BLI traces for b-P1aCy5-Cy3 RNA. Red: b-aminoP1a-Cy3 in 1 mM $MgCl_2$. Yellow: b-P1aCy5-Cy3 in 1 mM $MgCl_2$. Cyan: b-P1aCy5-Cy3 in 20 mM $MgCl_2$. Purple: no RNA.

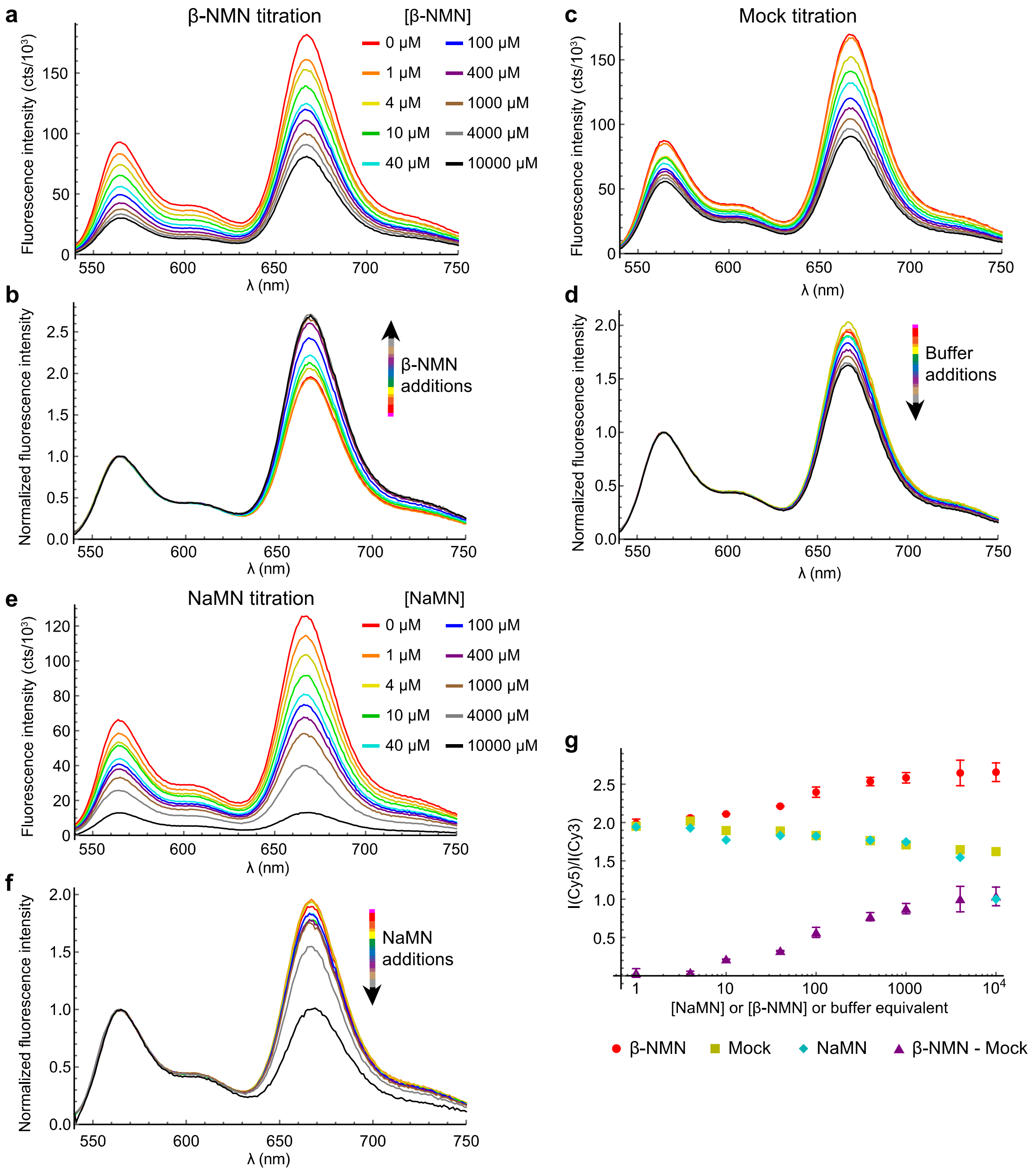


Figure S4. Bulk FRET measurements on b-P1aCy5-Cy3 RNA. (a) Raw fluorescence emission spectra following sequential additions of β-NMN to the indicated concentrations. (b) Spectra normalized to the peak of the Cy3 emission feature (reproduced from Fig. 3 for ease of comparison). (c-d) Raw (c) and normalized (d) spectra from a mock titration in which buffer was added in the same series of volume increments as β-NMN. Photobleaching causes a decrease in Cy5 emission relative to Cy3. (e-f) Raw (e) and normalized (f) spectra following sequential additions of nicotinic acid mononucleotide (NaMN) to the indicated concentrations. The trend closely tracks the mock titration until 4 mM NaMN. (g) Cy5 to Cy3 peak intensity ratio extracted from the β-NMN (red), mock (yellow) and NaMN (cyan) titration spectra. The photobleaching-corrected peak ratio used for fitting is plotted in purple.

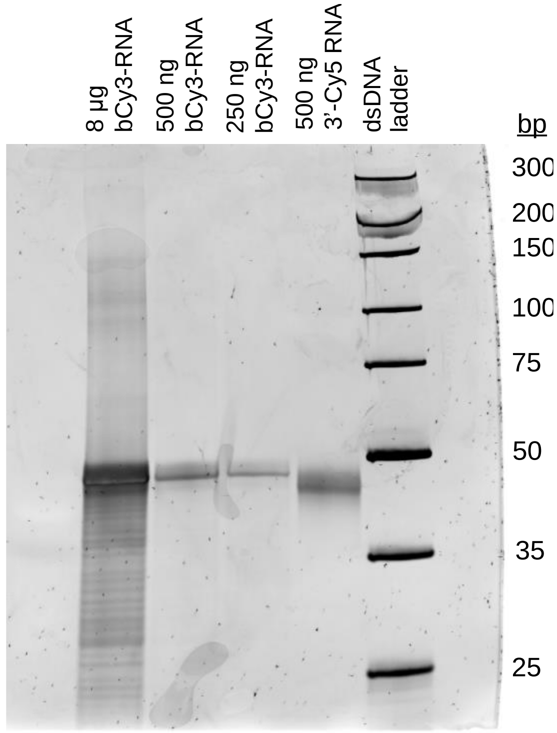


Figure S5. Raw image of gel in Fig. 4a.

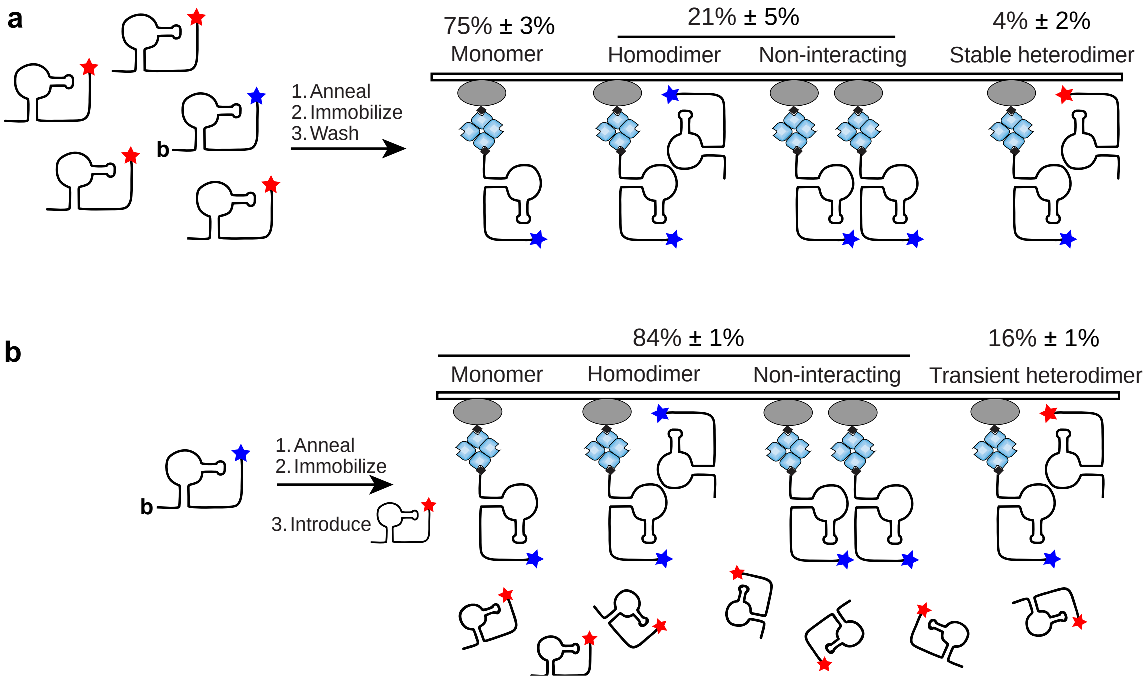


Figure S6. (a) Schematic of experimental design and results for “dimerization scenario 1”. 3’ Cy5 and b-aminoU14-Cy3 RNAs were annealed in a 10:1 ratio and immobilized on a microscope slide that was then thoroughly flushed with buffer. Imaging allowed the observation of monomers (75% of analyzed spots), stable homodimers and non-interacting colocalized molecules (21%) and stable heterodimers (4%). (b) Schematic of experimental design for “dimerization scenario 2”. b-aminoU14-Cy3 RNA was annealed and immobilized, then imaging was performed with 100 pM 3’ Cy5 RNA in the sample chamber. 16% of analyzed traces showed evidence of heterodimer formation with all colocalizations being transient.

### 3. Table of RNA sequences

| RNA | Sequence |
|---|---|
| b-aminoU14-Cy3 | 5'biotin-AUCUAUAGAGCGU(5-LC-N-U)GCGUCCGAAAGUCUAA ACAGACACGGCUCUUUAAAAACAAAAGGAGA-3'Cy3 |
| b-aminoP1a-Cy3 | 5'biotin-AUCUAUAGAGCGUUGCGUCCGAAAG(5-LC-N-U)CUAA ACAGACACGGCUCUUUAAAAACAAAAGGAGA-3'Cy3 |
| 3' Cy5 | AUCUAUAGAGCGUUGCGUCCGAAAGUCUAAACAGACACGGC UCUUUAAAAACAAAAGGAGA-3'Cy5 |
